# Growth factor-free, peptide-functionalized gelatin hydrogel promotes arteriogenesis and attenuates tissue damage in a murine model of critical limb ischemia

**DOI:** 10.1101/2023.05.24.542150

**Authors:** Corinne W. Curry, Sarah M. Sturgeon, Brian J. O’Grady, Alexis K. Yates, Andrew Kjar, Hayden A. Paige, Lucas S. Mowery, Ketaki A. Katdare, Riya V. Patel, Kate Mlouk, Madison R. Stiefbold, Sidney Vafaie-Partin, Atsuyuki Kawabata, Rachel M. McKee, Stephanie Moore- Lotridge, Adrienne Hawkes, Jiro Kusunose, Katherine N. Gibson-Corley, Jeffrey Schmeckpeper, Jonathan G. Schoenecker, Charles F. Caskey, Ethan S. Lippmann

**Author notes:** Correspondence to: Ethan S. Lippmann, Ph.D. 2400 Highland Ave 107 Olin Hall Nashville, TN 37212.

## Abstract

Critical limb ischemia (CLI) occurs when blood flow is restricted through the arteries, resulting in ulcers, necrosis, and chronic wounds in the downstream extremities. The development of collateral arterioles (i.e. arteriogenesis), either by remodeling of pre-existing vascular networks or *de novo* growth of new vessels, can prevent or reverse ischemic damage, but it remains challenging to stimulate collateral arteriole development in a therapeutic context. Here, we show that a gelatin-based hydrogel, devoid of growth factors or encapsulated cells, promotes arteriogenesis and attenuates tissue damage in a murine CLI model. The gelatin hydrogel is functionalized with a peptide derived from the extracellular epitope of Type 1 cadherins. Mechanistically, these “GelCad” hydrogels promote arteriogenesis by recruiting smooth muscle cells to vessel structures in both *ex vivo* and *in vivo* assays. In a murine femoral artery ligation model of CLI, delivery of *in situ* crosslinking GelCad hydrogels was sufficient to restore limb perfusion and maintain tissue health for 14 days, whereas mice treated with gelatin hydrogels had extensive necrosis and autoamputated within 7 days. A small cohort of mice receiving the GelCad hydrogels were aged out to 5 months and exhibited no decline in tissue quality, indicating durability of the collateral arteriole networks. Overall, given the simplicity and off-the-shelf format of the GelCad hydrogel platform, we suggest it could have utility for CLI treatment and potentially other indications that would benefit from arteriole development.

## Introduction

Critical limb ischemia (CLI) occurs when blood flow is restricted through the arteries, resulting in ulcers, necrosis, and chronic wounds in the peripheral extremities [1–3]. CLI is associated with several common comorbidities, including diabetes, obesity, and age, and the disease afflicts more than two million Americans. Given the complications associated with CLI, many patients do not qualify for invasive bypasses or stent placements that could reopen vasculature and restore blood flow to the limb [4–7]. As an alternate to these interventions, exercise therapy can naturally stimulate the development of collateral arterioles (i.e. arteriogenesis), which provides blood flow from distal sites to relieve tissue ischemia. However, many patients are non-compliant with exercise therapy, and “druggable” solutions to CLI do not exist. As a result, amputation is often the only viable path forward to prevent further complications, making CLI the leading cause of amputations in the United States [8]. This has led to a push for non-invasive therapeutic strategies that can promote arteriogenesis and restore blood flow as a treatment for CLI [9–11].

In clinical trials for CLI, early attempts were made to increase collateral vessel development by administering bolus injections of angiogenic growth factors, such as vascular endothelial growth factor (VEGF) and fibroblast growth factor (FGF), directly to the ischemic tissue [12–15]. These trials were based on positive outcomes in animal models [16,17]. In humans, these injections created vessel networks that generated short-term benefit, but lasting restoration of bulk flow was not achieved [12]. Since soluble growth factors are rapidly degraded or cleared from the site of injection, other trials have delivered plasmids or viral vectors to facilitate sustained production of growth factors by cells around the injury site, as well as different types of cells that can provide trophic support or contribute to new vessel formation [18–21]. However, these approaches have also failed to provide clinical benefit.

To address issues of growth factor clearance, researchers have more recently turned to hydrogels as scaffolds that enable sustained release of biologics [22,23]. Hydrogels are water-swollen networks composed of either natural or synthetic polymers, and their chemical and physical properties can be tuned for diverse biomedical applications [24]. Typically, hydrogels have been used as carriers of pro-angiogenic cargo for non-invasive CLI treatment, resulting in a number of biomaterial-biologic combinations in preclinical studies. For example, VEGF has been covalently bound or encapsulated in several hydrogel systems, and the prolonged release of VEGF to ischemic tissue increases vascular density, particularly in comparison to bolus delivery of VEGF without a supportive matrix [25–29]. Hydrogel delivery systems have also been used to encapsulate multiple cell types and growth factors in a single system [25,30–32]. However, these systems have not yet been transitioned to human trials, and the use of multiple biologics complicates routes to clinical translation.

To circumvent the need for encapsulated biologics, hydrogels have also been developed with conjugated peptides that mimic the activity of soluble proteins. This strategy reduces cost associated with recombinant proteins and can prolong biological activity since the bioactive cue is not depleted until the hydrogel degrades. Further, similar to how acellular ECM-based vascular grafts have shown great promise in bypass surgeries, peptide-functionalized hydrogels are more amenable to an “off-the-shelf” format that is compelling in a clinical setting. Several peptides have already been used for growth factor free vascularization strategies, for example a VEGF mimetic peptide that has been deployed in different hydrogel formats [33–35]. Indeed, self-assembling multidomain peptide hydrogels containing this particular peptide have been previously used to enhance perfusion in a CLI model [36]. Overall, these studies highlight the potential of using peptide-functionalized hydrogels for CLI treatment. However, this area of inquiry has not been extensively explored.

To augment these prior efforts, in this study, we sought to design a new peptide-functionalized hydrogel for CLI treatment. Since arterioles are believed to be the key vessel type for combatting tissue ischemia [37], we focused on mechanisms of arteriogenesis that could potentially be recapitulated with hydrogels. We used gelatin as the hydrogel backbone due to its low cost, ease of chemical modification, and natural ability to support endothelial cell growth [38]. Then, based on studies showing that N-cadherin-mediated signaling promotes smooth muscle cell (SMC) growth after vascular injury [39,40], we functionalized gelatin with a peptide from the extracellular domain of Type 1 cadherins, with the hypothesis that co-stimulation of endothelial cells and SMCs would better support arteriole assembly. This biomaterial, termed GelCad, was then crosslinked into a hydrogel *in situ* using a branched amine-reactive polyethylene glycol (PEG) polymer. Our study shows that GelCad hydrogels robustly promote collateral arteriole development *in vivo* and prevent ischemic damage in a severe femoral artery ligation model using mice with limited regenerative capacity. Overall, these outcomes highlight the promise of this simple, inexpensive, off-the-shelf hydrogel system for CLI treatment.

## Materials and Methods

### Ethics statement

All animal procedures were approved by the Vanderbilt Institutional Animal Care and Use Committee. Mice were housed in a 12-hour light/dark cycle and allowed ad libitum access to food and water.

### Gelatin functionalization strategy

10 grams of porcine skin gelatin (Thermo Fisher) were dissolved in 240 mL of phosphate buffered saline (PBS) for a concentration of 42 mg/mL. Hydrochloric acid (5M) was added dropwise until slightly acidic conditions were achieved (pH=6). Synthetic peptides were ordered from GenScript with a carboxylated N-terminus and acetylated C-terminus (N-Cadherin Sequence: Ac-HAVDIGGCE, Scrambled Sequence: Ac-AGVGDHIGCE). An intermediate reaction was used to prepare the peptides for binding. 80 milligrams of ethylcarbodiimide hydrochloride (EDC; Thermo Fisher) and 120 milligrams of n-hydroxysuccinimide (NHS; Thermo Fisher) were mixed with 80 milligrams of peptide. The combined solids were dissolved in 10 mL of N-N-dimethyformamide (DMF) (28 mg/mL), and 15 mL of PBS (19 mg/mL) at 37°C for 1 hour. The peptide intermediate was then added dropwise to the gelatin solution and reacted for 4 hours at 37°C with constant stirring. A second solution was prepared with 80 milligrams of 3-(4-hydroxyphenyl) propanoic acid (HPA; Sigma Aldrich), 80 milligrams EDC, and 120 milligrams NHS dissolved in 10 mL DMF and 15 mL PBS at 37°C for 1 hour; this solution was added dropwise to the gelatin-peptide solution and reacted at 37°C for 2 hours. Next, the reaction was quenched by raising the pH to 8 using sodium bicarbonate (5M). The final solutions were filtered using a tangential flow filtration system (10,000 kDa MW cutoff; Sartorius) with four filtration volumes of ultrapure water. After filtration, the biomaterial solutions were frozen, lyophilized, and pulverized for long term storage. Conjugation was confirmed with hydrogen nuclear magnetic resonance spectroscopy (^1^H-NMR). Specifically, the signature peaks for the peptide at 1.0 ppm and gelatin at 8.0 ppm were identified and the area under the curve was measured to determine the degree of functionalization.

### Hydrogel polymerization with microbial transglutaminase

For *ex vivo* experiments, gelatin-based biomaterials were crosslinked with microbial transglutaminase (mTG). Materials were prepared fresh for each experiment. A 10% (w/v) solution of biomaterial was prepared by manually mixing the pulverized biomaterial in Human endothelial serum free media (HESFM; Gibco) supplemented with fetal bovine serum (2% v/v) and PenStrep (1% v/v). The solution was fully dissolved after 30 minutes at 37°C. mTG was reconstituted in Hank’s Balanced Salt Solution (HBSS; Gibco) for a final concentration of 10% (w/v) and sterile filtered. To initiate polymerization, 35 μL of the mTG solution was used per milliliter of biomaterial and thoroughly mixed with a pipette.

### Hydrogel polymerization with polyethylene glycol

For *in vivo* experiments, gelatin-based biomaterials were crosslinked with a 4-arm PEG succinimidyl glutarate (20 kDa MW; JenKem). Gelatin-based biomaterials were reconstituted as 10% (w/v) solutions by stirring in sterile PBS (pH 6) at 37°C for thirty minutes. PEG was dissolved in sterile PBS (pH=6) to create a 10% (w/v) solution. To initiate polymerization, equal volumes of biomaterial and PEG solutions were thoroughly mixed with a pipette. Polymerization occurred within 90 seconds.

### Rheology

Bulk mechanical properties of the hydrogels were measured using a TA Instruments AR-G2 magnetic bearing rheometer. For hydrogels crosslinked with mTG, 600 μL of liquid was placed in PDMS molds (20 μm diameter) and allowed to solidify overnight before placing them on the rheometer disc. For hydrogels crosslinked with PEG, 600 μL of liquid was placed directly on the rheometer disc and allowed to solidify for 5 minutes. All moduli (G’ and G”) were determined using a logarithmic frequency sweep from 0.1 to 100 rad/s at 1% strain and 37°C.

### Atomic force microscopy

Local hydrogel stiffness (quantified as Young’s Modulus) was determined using a Bruker Dimension Icon atomic force microscope (AFM). All hydrogel samples were prepared as coverslip-bound discs for AFM measurements. Coverslips were glued to microscope slides and all measurements were performed in fluid at room temperature (approximately 25°C). All hydrogel measurements were taken using a pre-calibrated SAA-SPH-5UM AFM probe (Bruker) with a 0.2 N/m spring constant and 5-micron tip radius specifically designed for interfacing with soft biological samples. Additionally, at the start of every imaging session, a secondary calibration was completed using a 1 kPa polyacrylamide hydrogel standard as a reference for sample measurements. Three distinct 5 x 5 micron areas (256 individual measurements per area) were measured in tapping mode across three hydrogel samples for each condition, totaling nine captures per condition. The resulting force curves were baseline corrected, and Young’s modulus was calculated for each using the Hertzian model.

### Scanning electron microscopy

Microporous structures of the hydrogels were imaged using a Zeiss Merlin scanning electron microscope (SEM). Hydrogels were crosslinked as previously described, directly on the pin mount overnight. The next day, the samples were frozen and lyophilized. To expose the internal structure and flatten the topography, the dried hydrogel was cut to 1 millimeter in height with a razor. Samples were then gold coated and imaged. Pore diameter was calculated using ImageJ.

### *Ex vivo* arteriogenesis assay

Cortices from C57BL/6 female mice (2-4 months of age) were homogenized in ice cold HBSS. To remove debris, the homogenized samples were gently mixed with HBSS and centrifuged at 1,000 rpm for two minutes. After centrifuging, the debris and excess liquid were removed from the top of the micro centrifuge tube. The residual tissue was washed on a 50 μm mesh filter using 100 mL of PBS. Tissue remaining on the filter was divided into the prepared hydrogel conditions (100 μL of tissue/mL of hydrogel) and mixed with mTG at ratios described earlier. The tissue solution was then transferred to a glass bottom 24 well plate. HESFM (Gibco) supplemented with 1% PenStrep and 2% fetal bovine serum was added to the top of the crosslinked hydrogel after 3 hours. Medium was changed every three days. At designated time points, samples were fixed with 4% paraformaldehyde (PFA, Thermo Fisher) for one minute. After fixation, wells were washed 3 times with PBS for 15 minutes each. After washing, samples were blocked and permeabilized overnight in PBS containing normal goat serum (5%) and Triton-X (0.5%). Dy-Light Lectin dye at 647 (1:1000) and an anti-smooth muscle actin antibody (α-SMA, 1:500, Millipore, Cy3, mouse monoclonal) were then incubated with tissue for approximately 24 hours. Next, DAPI nuclear stain (1:1000) was added for at least 15 minutes. Samples were then imaged using a 40x lens with immersion oil on a Zeiss LSM 710 confocal microscope.

To quantify α-SMA coverage around lectin+ vessels in an unbiased manner, tube-like structures were randomly selected under the DAPI channel. Then, z-stacks were acquired across all fluorescence channels at 10-15 µm sections. Using the macros feature in Fiji, an ImageJ code was applied to all images to batch process and quantify the files. The stacked .czi files were opened in Fiji and projected into single channel images at maximum intensity. Each channel was contrast enhanced, despeckled, and rolling ball background subtracted at 100. In the lectin+ channel, a region of interest (ROI) was created around the vessel using the polygon tool and added to the ROI manager. The α-SMA channel was converted to a binary 8-bit image, and non-specific signal removed through an outlier subtraction and erosion. The ROI drawn from the lectin+ channel was then applied to the α-SMA and DAPI channels. Measurements of area, mean, and area fraction were used to establish binary signal for α-SMA on the lectin+ vasculature. DAPI+ nuclei were also counted using an identical batch process with an additional watershed and cellular analysis step.

### Injection and in situ crosslinking of hydrogels in the mammary fat pad

Female C57BL/6 mice (2-4 months old) were placed under anesthesia using isoflurane. A mixture of PEG-biomaterial solution was prepared as described above. Immediately after mixing, 150 µL of PEG-biomaterial solution was injected subcutaneously through a 16-gauge syringe into the mammary fat pad between the fourth and fifth nipples. The needle was removed after the hydrogel had fully solidified (approximately 1 minute). Each mouse received a gelatin hydrogel in one flank and a GelCad hydrogel in the other flank.

### Contrast enhanced ultrasound imaging of hydrogels in the mammary fat pad

Contrast enhanced ultrasound imaging was conducted on days 7 and 14 after hydrogel injections. Under anesthesia, mice received a 50 μL retro-orbital injection of microbubbles, and hydrogels were imaged using a research ultrasound system (Verasonics Vantage 256) with an L12-5 transducer driven at 9.2 MHz. Microbubbles were created in-house following established methods [41]. B-mode images were acquired at 500 Hz using plane wave imaging and filtered using singular-value decomposition filtering to isolate the microbubble signal, yielding Doppler flow images. For analyses, boundaries of the hydrogels were traced by hand in MATLAB using a combined B-mode and Power Doppler image to guide selections. Images were acquired and analyzed under fully blinded conditions.

### Immunohistological analyses of hydrogels in the mammary fat pad

Mice were cardiac perfused with 4% PFA. The hydrogels were harvested, and excess fat tissue was manually removed. Hydrogels were then fixed again in PFA overnight and stored in a 20% sucrose solution. For histology, hydrogels were embedded in optical cutting tissue medium (OCT; Tissue-Tek) and cryosectioned into 50 μm slices. Sections were cut every 500 μm along the frozen block for unbiased sampling. The collected sections were split in two groups for staining: 1) lectin (DyLight 649; 1:100; Thermo Fisher) and α-SMA (1:100; Millipore), and 2) CD31 (1:100; Abcam) and α-SMA. After overnight staining with primary antibodies, sections were counterstained for 1 hour at room temperature with secondary antibodies (donkey anti-rabbit Alexa Fluor 555 and donkey anti-mouse Alexa Fluor 647; 1:200; Thermo Fisher) and DAPI (1:1000; Thermo Fisher). Each section was imaged on a Zeiss 710 LSM confocal microscope with a 5x tile scan to obtain a complete overview of the sample. Arterioles within the hydrogel were identified by concentric rings of α-SMA, and additional images were acquired at higher magnification to confirm arteriole structure. For quantification of arteriole density, slices were selected at random without any identifiers and counted by a blinded observer.

### Femoral artery ligation model

Femoral artery ligations were performed on 8-12 week old BALB/c mice (Jackson Laboratories) as previously described [42]. Mice were anesthetized with isoflurane and placed in the supine position. The right femoral artery was located and separated from the femoral vein and nerve. Double knots were placed at the proximal position on the femoral artery, just above the lateral circumflex femoral artery. The distal location was identified on the femoral artery, just below the branch point for the popliteal artery, and two double knots were placed to fully prevent blood flow through the artery. 75 μL of PEG-biomaterial solution was then pipetted in the surgical sit, over the top of the ligated artery, and allowed to crosslink *in situ*. After the hydrogel solidified, the incision was closed with sutures. Laser doppler perfusion imaging (LDPI) was performed with a laser speckle perfusion imager (Perimed) immediately after the surgery and repeated 24 hours later to confirm success of the ligation, defined as reduced blood flow by at least 40% compared to the non-ligated limb. LDPI is described in more detail below.

### Assessment of tissue quality and ambulation after femoral artery ligation

Visual assessment of tissue quality and mobility were used to quantify ischemic severity on days 0, 1, 3, 7, and 14 post-ligation. Tissue quality scoring was based on a murine modified index with scores ranging from 0 (normal appearance) to 7 (complete limb loss).[43,44] Indicators between 0 and 7 were based on the amount of discoloration in the toenails, toes, and foot. The Tarlov mobility assessment was used to score movement of the ligated leg, with a score of 6 indicating fast rapid movements and 0 indicating no movement [45]. Scores between 0 and 7 were based on ability to bear weight, use of foot, and gait when moving.

### Laser doppler perfusion imaging analysis after femoral artery ligation

LDPI was performed on all animals at days 1, 3, 7, and 14 post-ligation to quantify perfusion in the ligated limb compared to the non-ligated control limb. While under isoflurane anesthesia, each mouse was placed on a heating pad at 37°C in the prone position with their shaved legs off the heating pad. Ten images were acquired and averaged per limb and the ROI drawn from the knee down. Relative perfusion was calculated as the ratio of signal in the ligated leg to the healthy leg.

### Assessment of histopathology after femoral artery ligation

On day 14 post-ligation, mice were cardiac perfused with 4% PFA fixative and transferred to the Vanderbilt Translational Pathology Shared Resource core facility for processing. Hindlimbs were collected and immersion fixed in 10% neutral buffered formalin for approximately 1 week, then decalcified in Immunocal (StatLab, McKinney, TX) for 48 hours. Tissues were routinely processed, embedded, cut at 4 μm thickness, and stained with hematoxylin and eosin. Tissue sections were then blindly evaluated and scored by a board certified veterinary pathologist. The following semi-quantitative scoring system was used to assess myodegeneration, myonecrosis, osteocyte loss (osteonecrosis) and bone marrow necrosis: 0 = not present; 1 = mild, rare, scattered pathology; 2 = moderate, multifocal marked, locally extensive pathology; 3 = diffuse pathology.

### Micro Computed Tomography imaging and quantification of vascular networks after femoral artery ligation

Mice were cardiac perfused with 4% PFA and subsequently with polymer casting liquid (MICROFIL®; Flow Tech Inc.) and left to solidify overnight. The mice were subsequently decalcified in EDTA (0.5 M) for five days, and Micro Computed Tomography (μCT) imaging was then performed on a Scanco microCT40 scanner. Image stacks were analyzed with Vesselucida® software, where user-guided and automatic directional kernels were used to trace the vessels in three dimensions. The reconstructed vessels were then visualized and measured in three dimensions. Data are presented as the ratio of the ligated limb to the healthy limb in the same mouse to account for variance between vascular casts.

## Results

### Synthesis and Characterization of Gelatin conjugated with a Cadherin Peptide (GelCad)

Previously, our group synthesized gelatin methacrylate conjugated with a peptide derived from the extracellular epitope of Type 1 cadherins [46]. This biomaterial was termed GelMA-Cad and could be polymerized into a hydrogel by photo crosslinking with UV light and a free radical initiator. Here, we updated our synthesis strategy to conjugate the peptide to gelatin using an EDC/NHS reaction. ^1^H-NMR was used to confirm presence of the peptides on gelatin through a triplet around 1.0 ppm, which is not present on gelatin alone (**Figure 1A**). We used a gelatin peak at 8.0 ppm as a reference and defined the degree of functionalization as the ratio of peptide to gelatin peaks; for the three batches of GelCad biomaterial synthesized in this study, the average degree of functionalization was 0.355±0.002, indicating excellent reproducibility. To move towards an *in situ* crosslinking formulation, we initially replaced the methacrylate group with 3-(4-hydroxyphenyl)propionic acid, which was intended to facilitate uniform polymerization with hydrogen peroxide, similar to previous descriptions [47]. However, this strategy yielded very soft hydrogels lacking mechanical integrity (data not shown), which we deemed unsuitable for translational work in sites where we needed better durability. Hence, we transitioned to polymerizing these “GelCad” hydrogels with either microbial transglutaminase (mTG, which crosslinks glutamine and lysine residues in the gelatin backbone) or a 4-arm polyethylene glycol succinimidyl glutarate (PEG, which reacts with free amines). mTG-crosslinked hydrogels were used for *ex vivo* experiments involving primary tissue because it rendered the hydrogels more translucent for immunofluorescent imaging, and PEG-crosslinked hydrogels were used for all *in vivo* experiments since PEG is a chemically defined reagent that is favorable for translational work.

**Figure 1:**
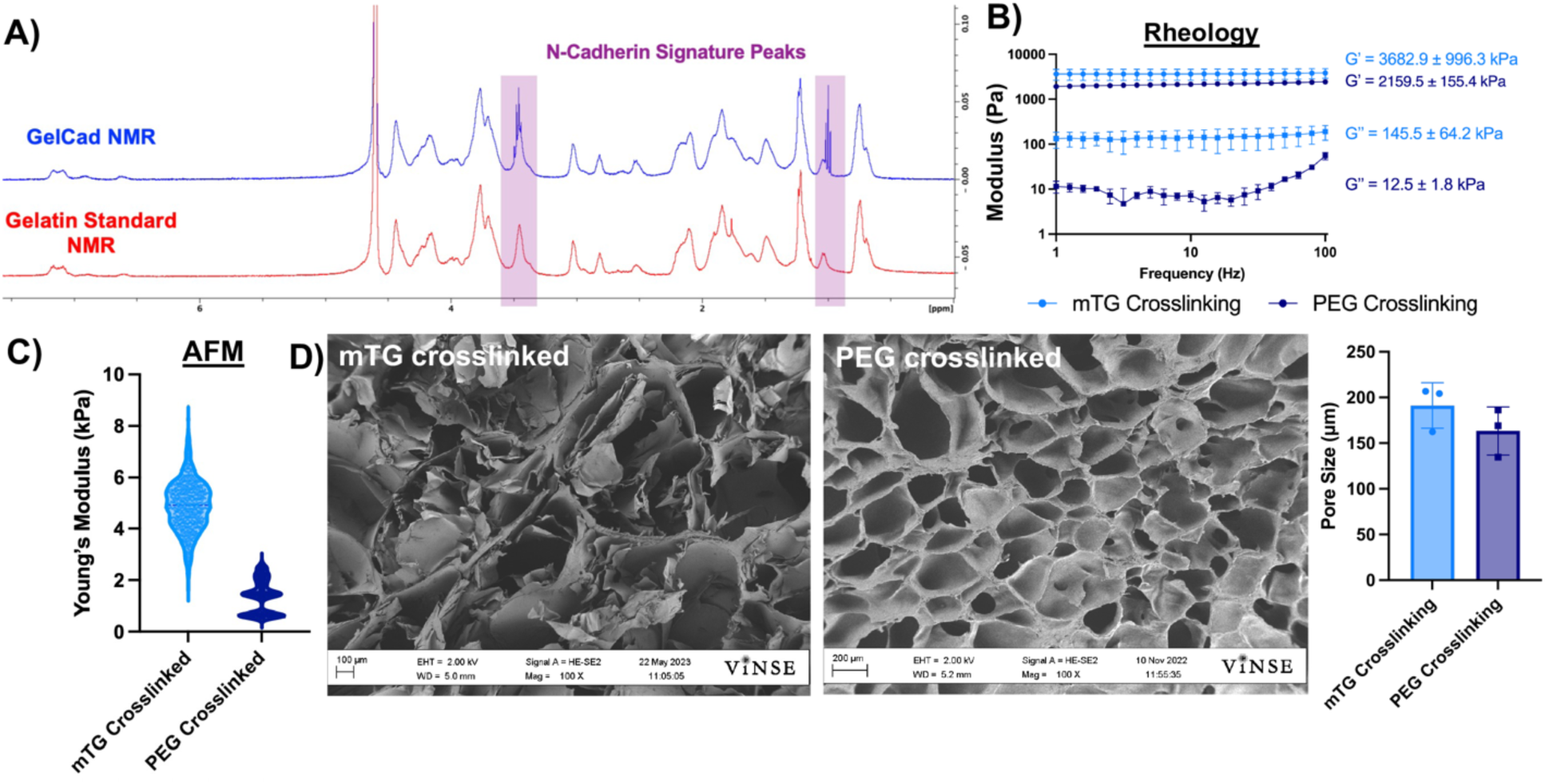
Characterization of GelCad hydrogels. **A)** Representative ^1^H-NMR of gelatin and GelCad samples. The shaded regions highlight the characteristic N-cadherin peptide peaks at 1.0 and 3.5 ppm. **B)** Rheology of mTG and PEG crosslinked GelCad hydrogels. Hydrogels were allowed to fully crosslink and then a frequency sweep was performed. Average storage and loss moduli are presented as a function of crosslinking method (mean ± SD from 3 independent replicates). **C)** AFM measurement of mTG and PEG crosslinked GelCad hydrogel stiffnesses. Data are presented as a violin plot of all points after outliers were removed (ROUT method at 10% aggression). Each data point represents a technical replicate. The average stiffness of mTG and PEG crosslinked hydrogels was 4.94 ± 1.8 kPa and 1.36 ± 0.95 kPa, respectively. **D)** Representative SEM images of mTG and PEG crosslinked GelCad hydrogels. For quantification of pore diameter, each data point represents the average diameter for a biological replicate that was calculated from 5 images per sample. Overall data are presented as mean ± SD from 3 biological replicates.

Both crosslinking strategies yielded soft, porous hydrogels as measured by rheology, atomic force microscopy, and scanning electron microscopy (**Figure 1B-E**). These formulations were transitioned to evaluation in *ex vivo* and *in vivo* assays.

### GelCad hydrogels promote recruitment of smooth muscle cells to vascular structures

Since prior work has shown that N-cadherin expression in SMCs is upregulated following vascular injury and modulates SMC proliferation [39], we hypothesized that the presence of a cadherin-mimicking peptide in gelatin would enhance SMC recruitment to vessel structures and thereby facilitate arteriole formation. To assess this possibility, we homogenized primary mouse brain cortices to expose vessels and simulate injury conditions. We then embedded this *ex vivo* tissue into mTG-crosslinked gelatin, GelCad, or “GelScram” hydrogels (where the latter contains a scrambled version of the cadherin peptide). As indicated by ⍺-SMA signal, tissue embedded in GelCad hydrogels had significantly higher coverage of SMCs on lectin+ vessel-like structures after 24 hours relative to either control (**Figure 2**). After 7 days, vascular SMC coverage remained significantly higher in GelCad hydrogels relative to gelatin, while SMC coverage in GelScram hydrogels increased to levels that were statistically insignificant with respect to GelCad hydrogels (**Figure 2**). We believe the narrowed difference is due to latent biologic activity of the scrambled peptide. Qualitatively, we also note that the SMCs in GelCad hydrogels formed concentric circles around the vessel structures, typical of arterioles, while SMCs in GelScram hydrogels remained more disorganized. Overall, these outcomes motivated further investigation of GelCad hydrogel performance *in vivo*.

**Figure 2:**
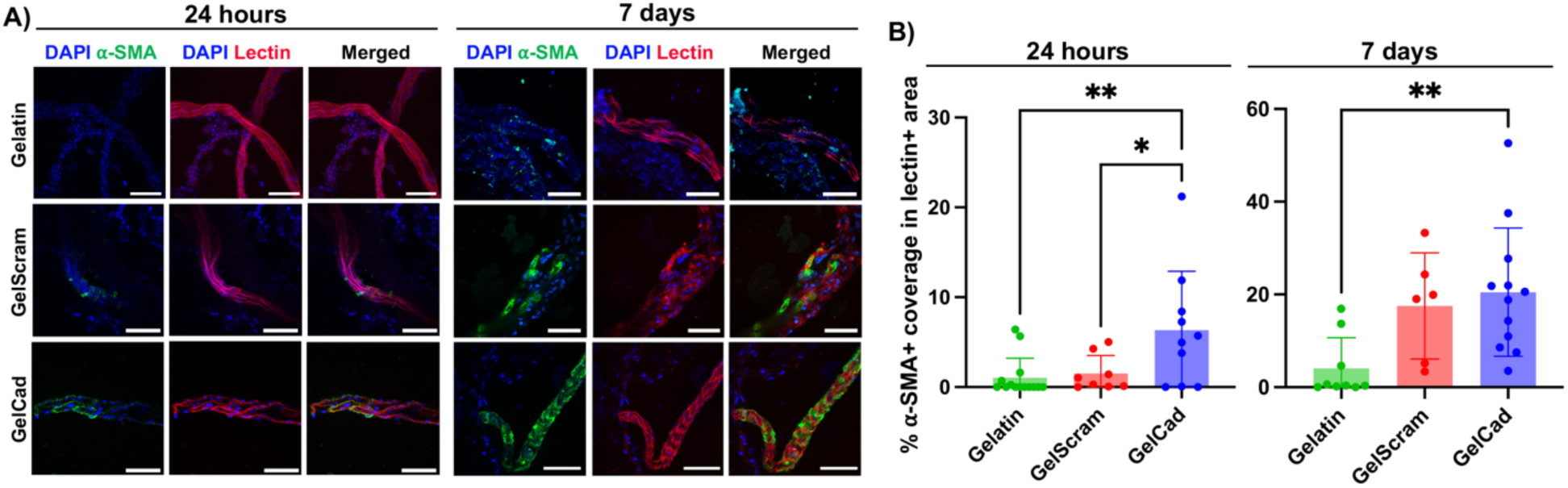
Development of arterioles in primary tissue embedded in GelCad hydrogels. **A)** Representative images of ⍺-SMA coverage on lectin+ vessel-like structures 24 hours and 7 days after embedding of primary mouse cortical brain tissue. Scale bars, 50 μm. **B)** Quantification of ⍺-SMA coverage on lectin+ vessel-like structures 24 hours and 7 days after embedding. Each data point represents an individual image (N=6-14 biological replicates). Statistical significance at each time point was calculated by one-way ANOVA with Dunnett’s multiple comparison test (*, p<0.05; **, p<0.01). The image processing strategy used for quantification is shown in **Supplemental Figure 1**.

### Evaluation of arteriole development in GelCad hydrogels after injection into the mammary fat pad

To build on our *ex vivo* outcomes, we next examined the performance of PEG-crosslinked hydrogels in the mammary fat pads of C57BL/6 mice. This site was chosen based on ease of *in situ* evaluation and recovery of hydrogels for postmortem analyses. After hydrogel injections, we first examined vascular development using contrast enhanced ultrasound. At day 7 post-injection, large vessels could be detected on the interior of gelatin and GelCad hydrogels, with potentially more vessels in the GelCad samples (**Figure 3A**). However, it was difficult to quantify vessel features because the absolute margin between the hydrogel and surrounding tissue had to be estimated and manually drawn. To address this issue, at day 14, we isolated the intact hydrogels for cryosectioning and immunohistochemistry. In GelCad hydrogels, we could readily locate numerous α-SMA+ vessels within the hydrogel indicative of arteriole formation, whereas in gelatin hydrogels, we could locate some putative arterioles but the α-SMA signal intensity was much lower (**Figure 3B**); quantification of total arteriole numbers revealed significant differences (**Figure 3C**), which mirrored our *ex vivo* data in **Figure 2**. Collectively, these results further indicate that the GelCad hydrogels enhance arteriole development.

**Figure 3:**
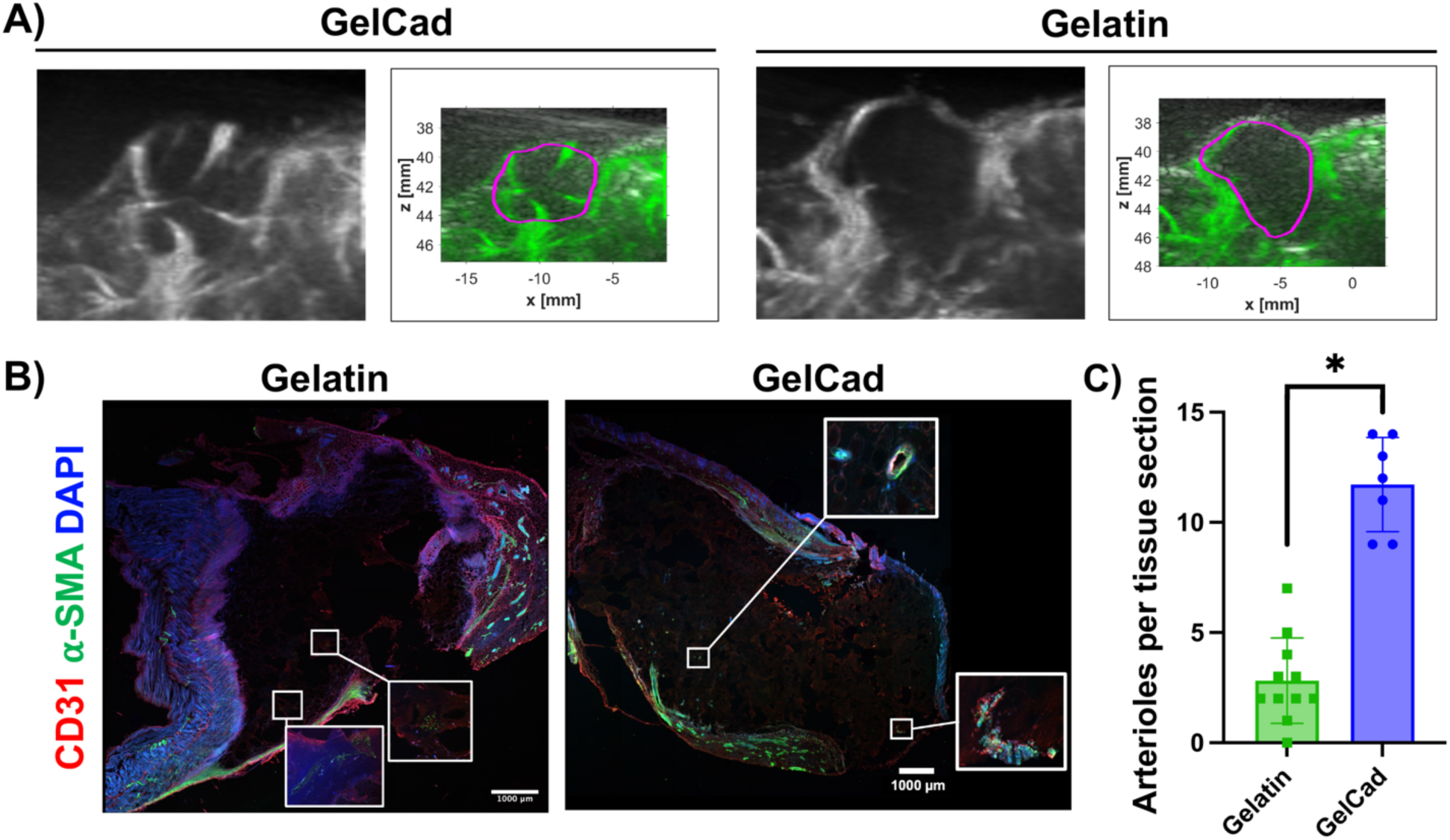
Evaluation of arteriole development in GelCad hydrogels after injection in mouse fat pad. **A)** Representative images of contrast enhanced ultrasound at day 7. The black and white B-mode images show the boundaries of the mouse body, and the Power Doppler images in green highlight the vasculature. The purple boundary marks the estimated boundaries of the hydrogel. The full set of images can be found in **Supplemental Figure 2**. **B)** Representative images of cryosectioned hydrogels that were extracted at day 14. The insets provide higher magnification of areas with ⍺-SMA to highlight weaker, more diffuse signal in gelatin relative to GelCad hydrogels. **C)** Quantification of arterioles per tissue section in each condition. Each data point represents an individual tissue section, and data are presented as mean ± SD from N=2-3 biological replicates per condition. Statistical significance was calculated from a student’s paired t-test (*, p<0.05).

### Evaluation of GelCad hydrogel efficacy in a mouse model of critical limb ischemia

Next, we sought to evaluate the therapeutic potential of the GelCad hydrogel platform in a femoral artery ligation model of CLI, with the hypothesis that its arteriogenic properties would enhance reperfusion and prevent ischemic deficits. For these experiments, we chose to use BALB/c mice because this strain has the lowest density of endogenous collateral vessels and exhibits poor recovery after femoral artery ligation [48,49], suggesting that any benefits facilitated by the hydrogels would be easy to delineate. To carry out this model, we tied double knots around two distal locations on the femoral artery in the hindlimb to create a severe ischemic event, then crosslinked hydrogels over the tissue and closed the surgical site. We used LDPI to confirm success of the ligation as indicated by blood flow reductions greater than 60%. Over 14 days, we evaluated recovery of blood flow, mobility, and tissue quality in mice treated with gelatin, GelScram, or GelCad hydrogels, as well as a sham control that received saline. Using a modified ischemia index, mice treated with GelCad hydrogels had significantly less necrosis in the ligated limb compared to all other conditions; of note, mice receiving gelatin hydrogels or saline typically lost their limb due to severe necrosis between days seven and fourteen, highlighting the severity of this model (**Figure 4A-B**). Using the Tarlov mobility index, we also determined that mice receiving GelCad hydrogels had near perfect mobility restored after 7 days, whereas mice receiving gelatin hydrogels or saline never regained mobility (**Figure 4C**).

**Figure 4:**
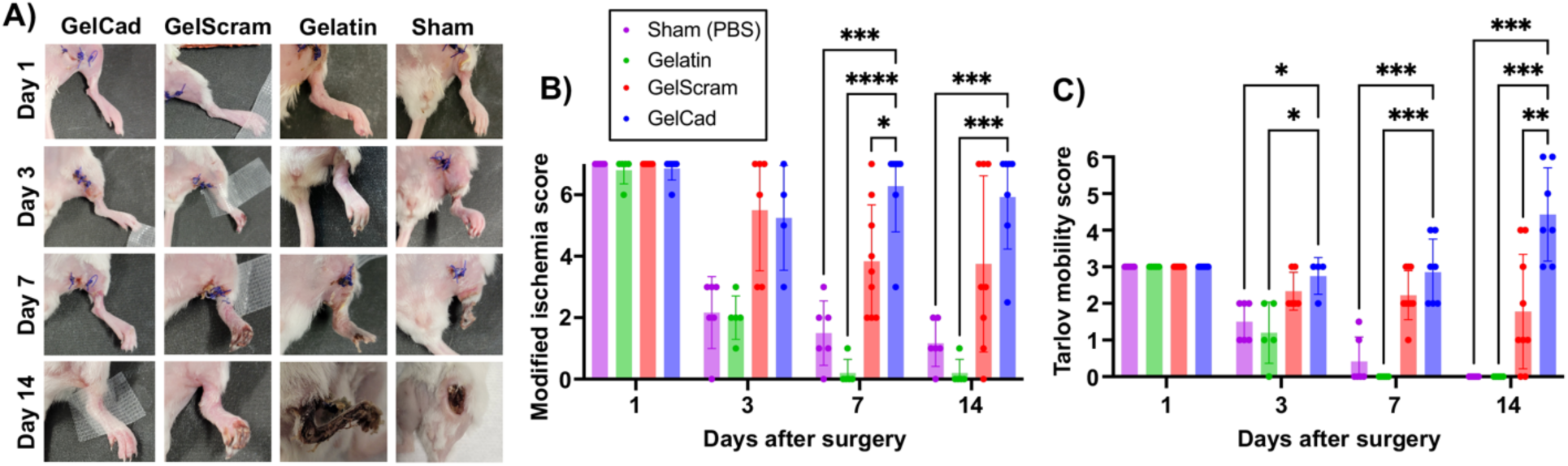
Physical appearance and mobility of mice after femoral artery ligation and hydrogel delivery. **A)** Representative images of the ligated hindlimb at each assayed time point as a function of treatment. **B)** Modified ischemia scores (0-7 scale) at each assayed time point as a function of treatment. Each data point represents an individual mouse and data are presented as mean ± SD from N=5-9 biological replicates per condition. A two-way ANOVA mixed effects model with the Geisser-Greenhouse correction and Dunnett’s multiple comparison test was used to assess statistical significance (*, p<0.05; ***, p<0.001; ****, p<0.0001). **C)** Tarlov mobility scores (0-6 scale) at each assayed time point as a function of treatment. Each data point represents an individual mouse and data are presented as mean ± SD from N=5-9 biological replicates per condition. A two-way ANOVA mixed effects model with the Geisser-Greenhouse correction and Dunnett’s multiple comparison test was used to assess statistical significance (*, p<0.05; **, p<0.01; ***, p<0.001).

These outcomes mirrored the LDPI measurements, where mice receiving GelCad hydrogels had a near perfect restoration of perfusion (1:1 ratio to the healthy leg) within 14 days and mice treated with gelatin hydrogels or saline did not exhibit restored perfusion (**Figure 5A-B**). Of interest, mice treated with GelScram hydrogels exhibited very heterogeneous responses, which likely reflects the latent biological activity observed in the *ex vivo* arteriole sprouting assay. In some cases, these mice recovered well, whereas in others the limb became necrotic, in contrast to the GelCad hydrogels that consistently prevented ischemic tissue damage. To further confirm collateral development, we perfused mice with a radiopaque polymer and performed μCT imaging according to established techniques [50]. Then, we used Vesselucida software to trace and quantify the vascular networks in healthy limbs versus ligated limbs treated with GelCad hydrogels. Of note, despite the ligated limb lacking blood flow through the femoral artery, total vessel length in the ligated limb was roughly equivalent to the healthy limb, and no significant differences in vessel diameter were noted (**Figure 5C-E**)—these data suggest compensation of blood flow through increased collateral density.

**Figure 5:**
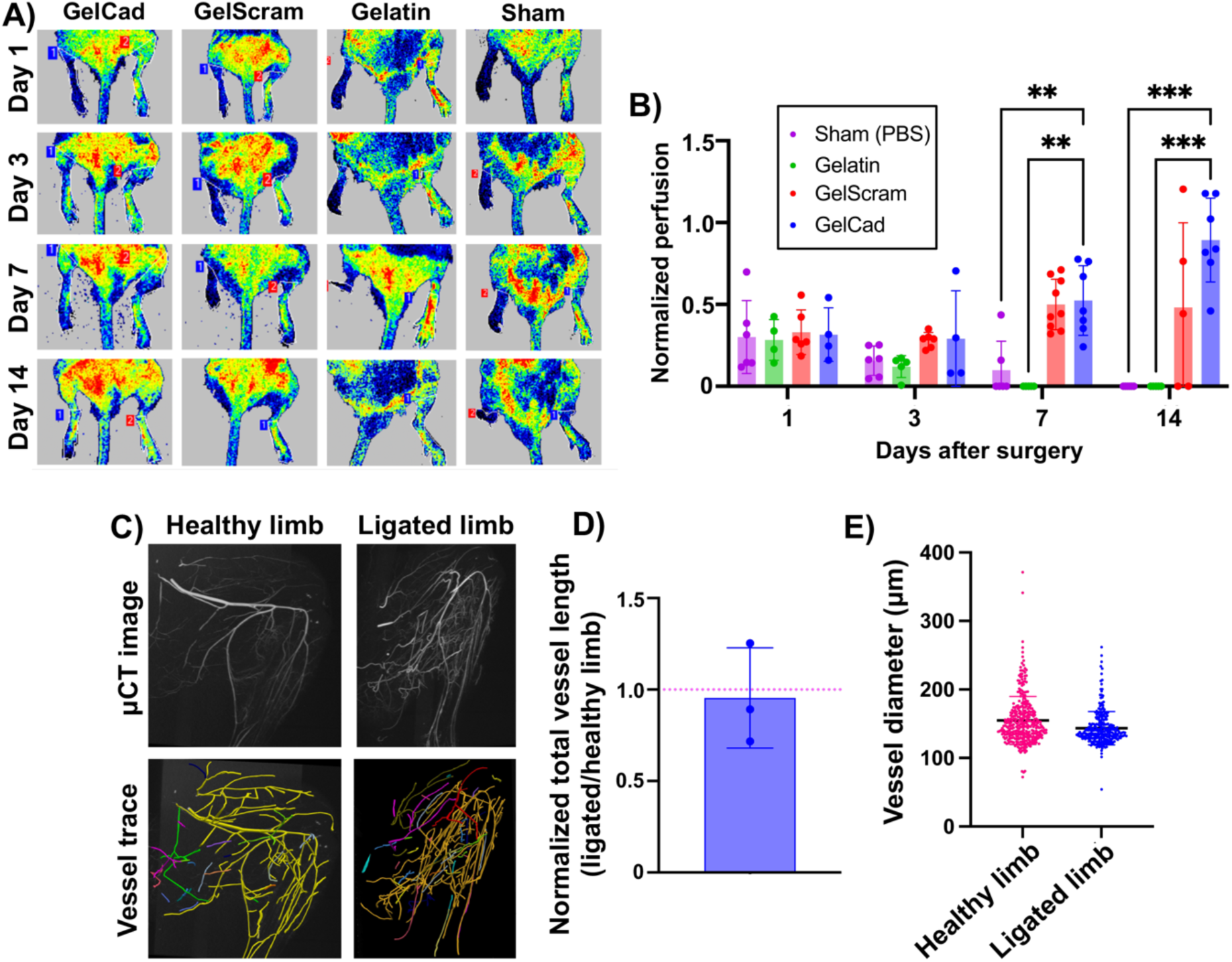
Recovery of perfusion after femoral artery ligation and hydrogel delivery. **A)** Representative LDPI images at each assayed time point as a function of treatment. For each mouse, healthy limbs are on the right side and ligated limbs are on the left side. **B)** Quantification of normalized perfusion (ratio of LDPI signal in ligated limb relative to healthy limb in the same mouse) as a function of treatment at each assayed time point. Each data point represents an individual mouse and data are presented as mean ± SD from N=5-9 biological replicates per condition. A two-way ANOVA mixed effects model with the Geisser-Greenhouse correction and Dunnett’s multiple comparison test was used to assess statistical significance (**, p<0.01; ***, p<0.001). **C)** Representative μCT images and corresponding vessel traces of healthy versus ligated limb in a mouse treated with a GelCad hydrogel. In the vessel trace images, colors indicate annotation of individual vessel networks. Colors are automatically and randomly assigned by the software to visually separate the vascular networks when tracing. The workflow for this imaging analysis pipeline is shown in **Supplemental Figure 3**. **D)** Quantification of total vessel length in the hindlimbs. To account for variance in perfusion with the MICROFIL polymer, data are presented as normalized vessel lengths for ligated versus healthy hindlimb in the same mouse. Each data point represents an individual mouse and data are presented as mean ± SD from N=3 biological replicates. The red line indicates the expected ratio for two healthy limbs. **E)** Aggregated vessel diameters from the vessel trace data for healthy and ligated limbs. Each datapoint represents a single vessel (determined from branch point analyses). Data are presented as mean ± SD from N=3 biological replicates. Statistical significance was calculated by a student’s paired t-test (no significant difference).

Finally, to confirm that restored blood flow had significant effects on physiology beyond visual appearance in **Figure 4**, a board-certified veterinary pathologist performed a blinded histopathologic evaluation of myodegeneration, myonecrosis, osteocyte loss, and bone marrow necrosis using semi-quantitative metrics (**Figure 6**). Here, mice treated with GelCad hydrogels had significantly less severe muscle degradation and necrosis of muscle and bone marrow relative to all other conditions. Of note, necrosis was almost completely absent in mice treated with GelCad hydrogels but pervasive in all other conditions. Further, no osteocyte loss was observed in mice treated with GelCad hydrogels versus some mild loss in the other conditions. Hence, GelCad hydrogels prevent tissue degeneration caused by ischemic injury, likely by rapidly restoring blood flow through collateral development.

**Figure 6:**
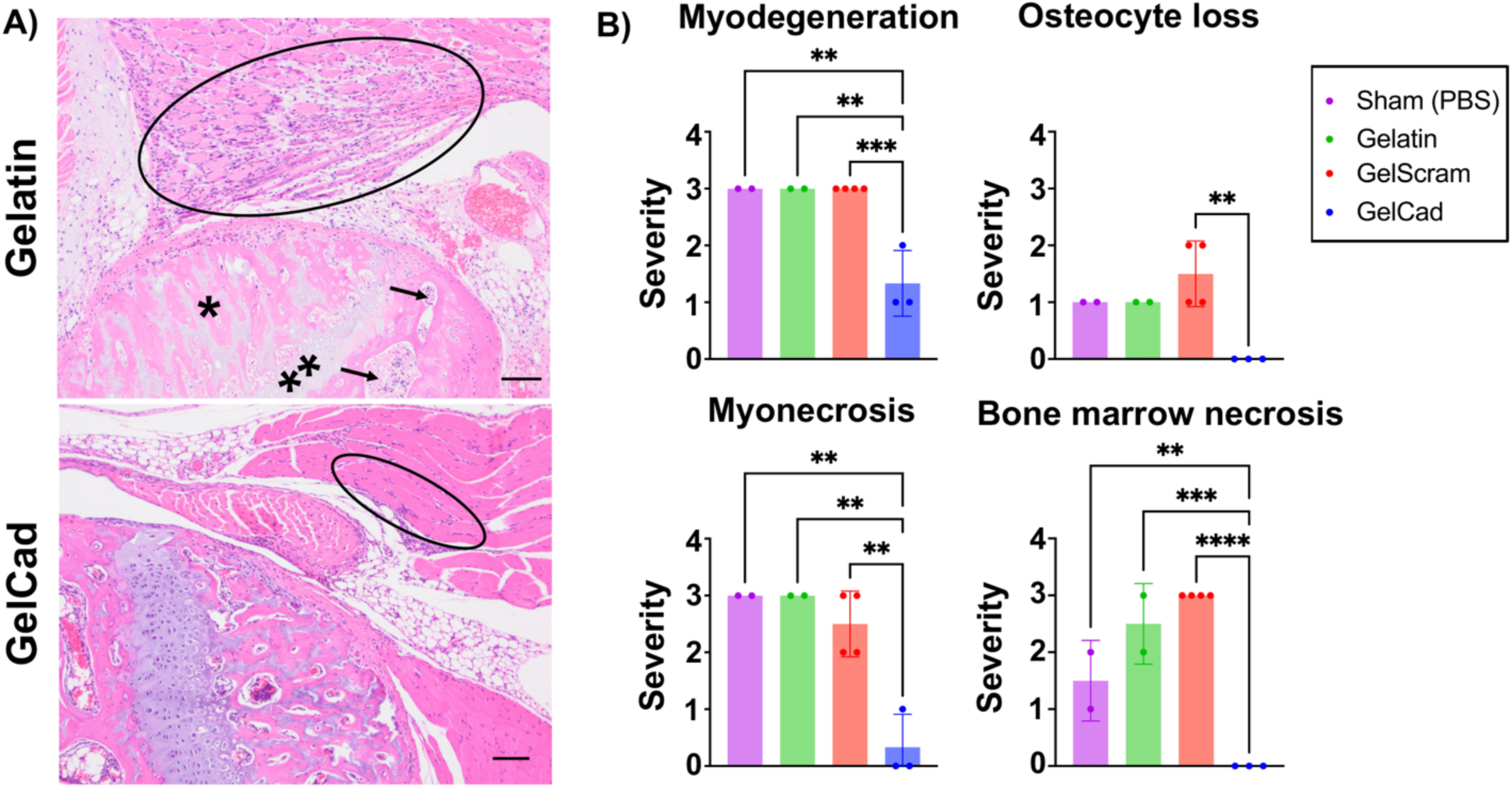
Histopathology of ligated hindlimbs receiving different hydrogel treatments. **A)** Representative photomicrographs of H&E-stained images of the knee in ligated limbs of mice treated with gelatin versus GelCad hydrogels. In the gelatin image, marked myoregeneration with scattered myodegeneration and necrosis is encircled, and severe necrosis of the bone marrow (arrows), osteonecrosis (*), and chondronecrosis (**) are evident. In the GelCad image, only very mild myoregeneration is noted (encircled). **B)** Quantification of pathology (0-3 scale). GelCad and GelScram samples were evaluated at day 14 post-ligation, and gelatin and sham samples were evaluated at day 7 post-ligation. Each data point represents the score from a single mouse. Data are presented as mean ± SD from N=2-4 biological replicates per condition. Statistical significance was calculated using a one-way ANOVA with Dunnett’s multiple comparisons test (**, p<0.01; ***, p<0.001; ****, p<0.0001).

Bolstered by these positive results, we selected two mice who received the femoral artery ligation and treatment with GelCad hydrogels and aged them for 5 months to assess long-term durability of therapeutic responses. At the endpoint, mice had normal ambulation and no visible signs of necrosis in the ligated limbs (**Figure 7A**). LDPI imaging indicated a near 1:1 perfusion ratio between the ligated and healthy limbs, indicating robustness of the collateral network and continued compensation for diminished circulation from the femoral artery (**Figure 7B-C**). Curiously, tracing of the collateral networks in μCT images revealed potential increases in total vessel length but not vessel diameter, which potentially suggests growth of new collateral vessels at this extended time point (**Figure 7D-F**). However, the small sample size prevents any definitive conclusions. Since we know that GelCad hydrogels produce this durable response, future work will focus on analyzing these extended time points to better understand long-term mechanisms of collateral development as a function of these hydrogels.

**Figure 7:**
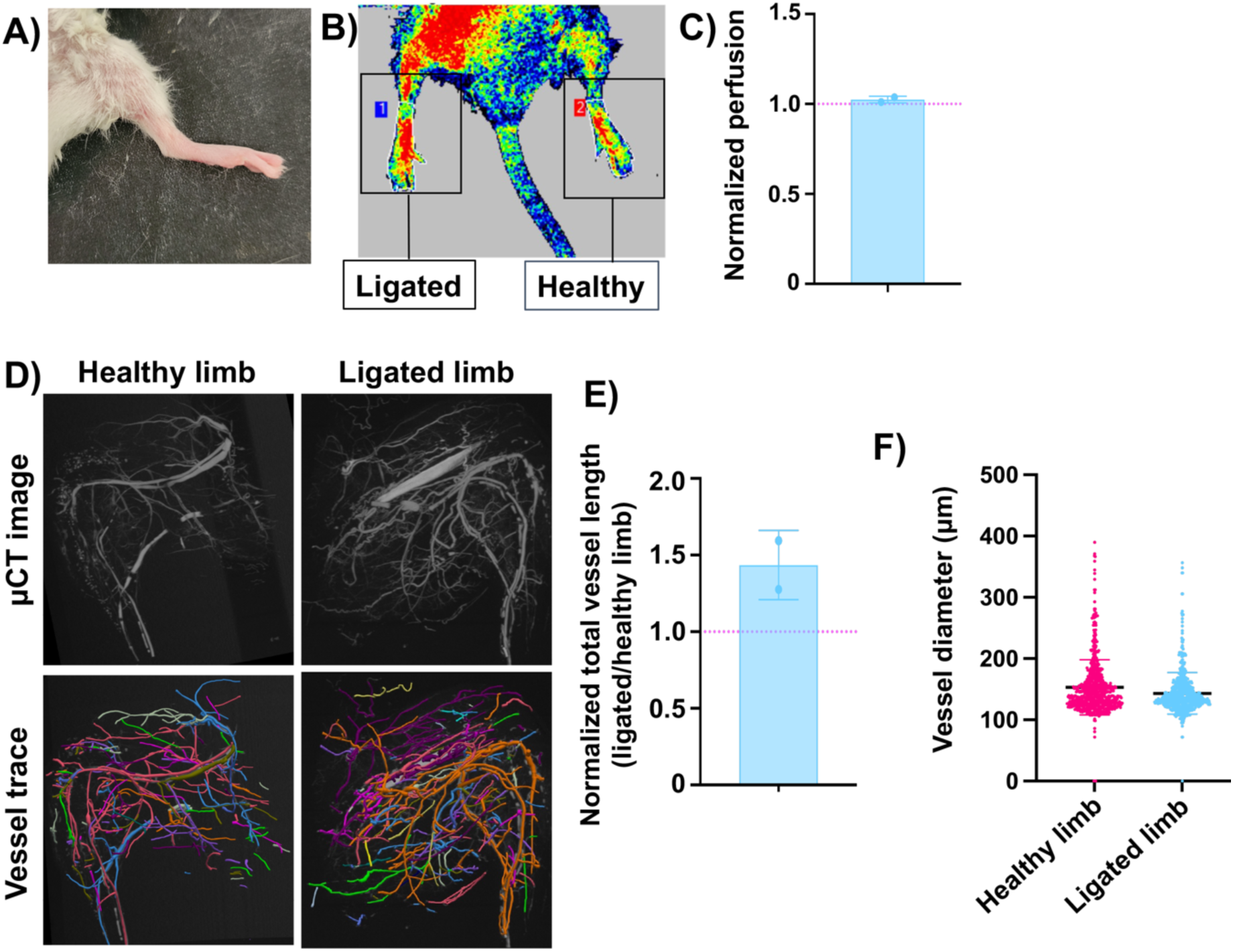
Evaluation of mice 5 months after femoral artery ligation and treatment with GelCad hydrogels. **A)** Representative image of hindlimb appearance. **B-C)** Representative image and quantification of LDPI measurements. Data are presented as mean ± SD from N=2 biological replicates. The red line indicates the expected ratio for two healthy limbs. **D)** Representative μCT images and corresponding vessel traces of a healthy versus ligated limb. **E)** Quantification of total vessel length in the hindlimbs. To account for variance in perfusion with the MICROFIL polymer, data are presented as normalized vessel lengths for ligated versus healthy hindlimb in the same mouse. Each data point represents an individual mouse and data are presented as mean ± SD from N=2 biological replicates. The red line indicates the expected ratio for two healthy limbs. **F)** Aggregated vessel diameters from the vessel trace data for healthy and ligated limbs. Each datapoint represents a single vessel (determined from branch point analyses). Data are presented as mean ± SD from N=2 biological replicates. Statistical significance was calculated by a student’s paired t-test (no significant difference).

## Discussion

In this study, we evaluated the arteriogenic activity of cadherin peptide-functionalized gelatin hydrogels. We determined that the peptide played a key role in recruiting SMCs to vessels, which facilitated arteriole development *ex vivo* and *in vivo*. In a severe CLI model, GelCad hydrogels contributed to blood flow restoration and prevented ischemic tissue damage over many months. Overall, our data suggest GelCad hydrogels have promising therapeutic potential in CLI and other conditions where development of collateral arterioles would be beneficial.

With respect to other hydrogels, the main benefit of the GelCad platform is the bioactivity achieved without release of soluble growth factors. Pioneering work by Dr. David Mooney and colleagues showed that sustained release of both vascular endothelial growth factor VEGF and platelet-derived growth factor (PDGF) from poly(lactide-co-glycolide) scaffolds could promote the formation of dense blood vessels lined with ⍺-SMA+ SMCs [51]. More recent approaches have leveraged alginate for growth factor release, for example VEGF and insulin-like growth factor delivery in mouse and rabbit hindlimb ischemia models [31]. In contrast to these passive growth factor release strategies, other hydrogel platforms have been engineered for active release of growth factors to promote vascularization, for example ultrasound-mediated release of bFGF from fibrin-based scaffolds [52,53]. While many of these approaches have demonstrated therapeutic benefit in CLI models, their clinical translation could be hampered by the difficulty and expense of producing recombinant growth factors at scale under GMP conditions. The GelCad hydrogels, consisting solely of gelatin and a short synthetic peptide, are much easier and cheaper to synthesize and store as an “off-the-shelf” therapy. The use of gelatin as an inexpensive and biocompatible scaffold also lends well to translational initiatives.

To evaluate the efficacy of GelCad hydrogels, we intentionally chose the BALB/c mouse strain with limited capacity for endogenous recovery from hindlimb ischemia; since most studies use strains with better recovery, such as C57BL/6 or FVB, direct comparisons to most other published outcomes is not possible. However, we note two recent studies in BALB/c mice that can be used as reference points. The first study used the aforementioned acoustically-responsive scaffolds for bFGF release, which improved the ischemia score above the negative controls, but mice receiving no intervention only had to be euthanized ∼20% of the time due to excessive necrosis [52]. This contrasts our study where all of our negative control mice autoamputated, suggesting our injury model was much more severe and again highlighting the efficacy of the GelCad hydrogels. A second study used gelatin hydrogels synthesized as bulk constructs or containing microchannels to encourage blood vessel infiltration [54]. Here, 100% of mice receiving saline or a bulk gelatin hydrogel after femoral artery ligation exhibited some degree of toe, foot, or limb loss. Yet, in mice receiving the microchannel-laden hydrogels, >60% of mice still lost a foot or toe. Again, this contrasts our study where mice receiving GelCad hydrogels had no visible evidence of tissue damage, let alone appendage loss. Further, in this study, microchannels had to be pre-formed in the hydrogels, which were then implanted at the injury site; this extra step would potentially complicate delivery in a clinical setting, whereas our injectable GelCad hydrogel formulation does not have this concern.

A key limitation of this current study is that we have not determined how cadherin peptide density influences arteriole development. Our current formulation is based on our prior work on neural tissue models, which also did not explore the variable of peptide density. Signaling through cadherins can activate many extracellular cascades such as, β-catenin and Rho family GTPases, and we suspect the combined activation of cadherin signaling by the synthetic peptide and integrin signaling by the gelatin backbone underlies arteriole assembly [55–59]. It is possible that altering peptide density could further improve this phenomenon, and we plan to explore this design variable in future studies. We also note intriguing possibilities for further improving GelCad hydrogel performance, such as tethering additional peptides that mimic the activity of soluble growth factors or transitioning from a bulk hydrogel formulation to granular hydrogel microparticles [33,34,60,61]. Crucially, each of these strategies would still allow for growth factor-free formulation and injectable delivery, thus maintaining key advantages over other hydrogel platforms used in CLI. While the first-generation GelCad platform shows excellent efficacy in a mouse model, larger species may require a higher density of collateral arterioles to achieve therapeutic benefit, thus motivating these proposed improvements to the GelCad formulation. In addition, in future work, it will be important to benchmark the efficacy of the GelCad platform in models that superimpose CLI co-morbidities like age, obesity, and diabetes, since our work and most other studies solely utilize young and healthy mice. Overall, the initial performance of the first-generation GelCad hydrogel system and its capacity for further modification highlights its promise for future clinical intervention.

## Author contributions

CWC and ESL conceived the project structure with input from SMS and BJO. CWC performed the majority of experiments and all data analyses. SMS developed the strategies for hydrogel injections into mouse fat pads and BJO developed the hydrogel synthesis strategy and the experimental framework for the *ex vivo* assays. AY performed atomic force microscopy. AKjar assisted with hydrogel imaging. LSM performed rheology. MRS assisted with hydrogel syntheses. SV-P assisted with histology and blinded arteriole counting. HAP, RVP, KM, and KAK assisted with animal experiments and hydrogel characterization. AH, JK, and CFC performed and analyzed ultrasound experiments. AKawabata, RMM, SM-L, and JGS assisted with MICROFIL perfusion and μCT imaging. KNG-C performed blinded histopathology assessments. JS performed femoral artery ligations.

## Acknowledgments

Primary funding for this work was provided by a Chan Zuckerberg Initiative Ben Barres Early Career Acceleration Award (grant 2019-191850 to ESL) and the NIH (grant R01 NS110665 to ESL). CWC and AK were supported by the National Science Foundation Graduate Research Fellowship Program. BJO and AY were supported by the Vanderbilt Interdisciplinary Training Program in Alzheimer’s Disease (NIH grant T32 AG058524). Support for AFM and SEM was provided by the Vanderbilt Institute of Nanoscale Science and Engineering. NMR was conducted in the Vanderbilt Biomolecular NMR facility, which is supported in part by grant 0922862 from the NSF and grant S10 RR025677 from the NIH. Histology was conducted by the Vanderbilt Translational Pathology Shared Resource core facility, which is supported in part by NIH grant P30 CA068485. Imaging was conducted in the Vanderbilt Cell Imaging Shared Resource facility, which is supported in part NIH grants P30 CA068485, P30 DK058404, and P30 EY008126. The authors would like to thank Dr. Leon Bellan for use of the TA Instruments rheometer and Zeiss LSM 710 confocal microscope and Dr. Craig Duvall for use of the Perimed laser speckle perfusion imager.

