## Supplemental information for "Growth factor-free, peptide-functionalized gelatin hydrogel promotes arteriogenesis and attenuates tissue damage in a murine model of critical limb ischemia"

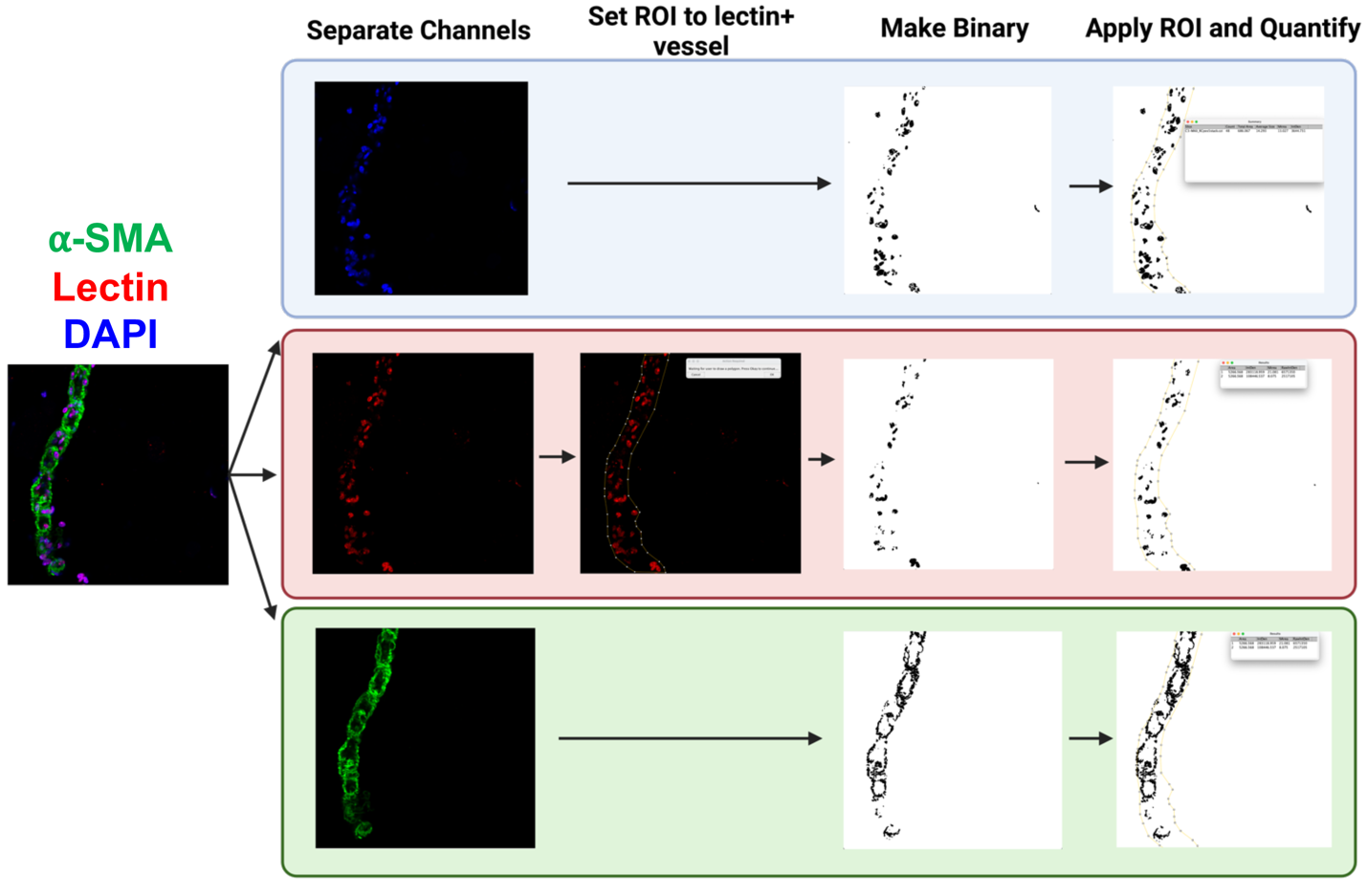

**Supplemental Figure 1: Example of image analysis pipeline used to quantify SMC coverage on vessel-like structures in *ex vivo* tissue samples embedded in hydrogels.**

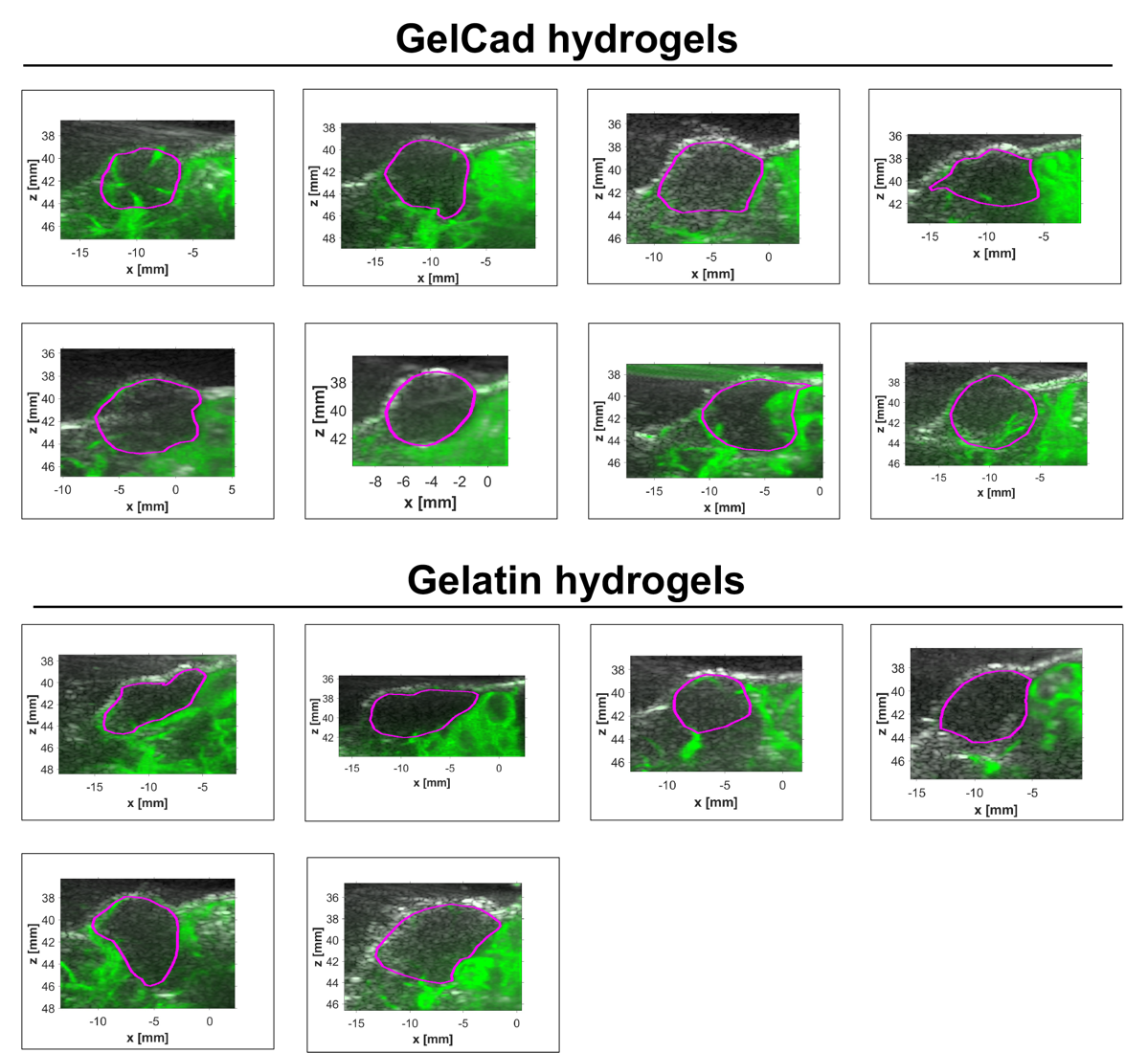

**Supplemental Figure 2: Full set of contrast enhanced ultrasound images of hydrogels in the mouse fat pad.**

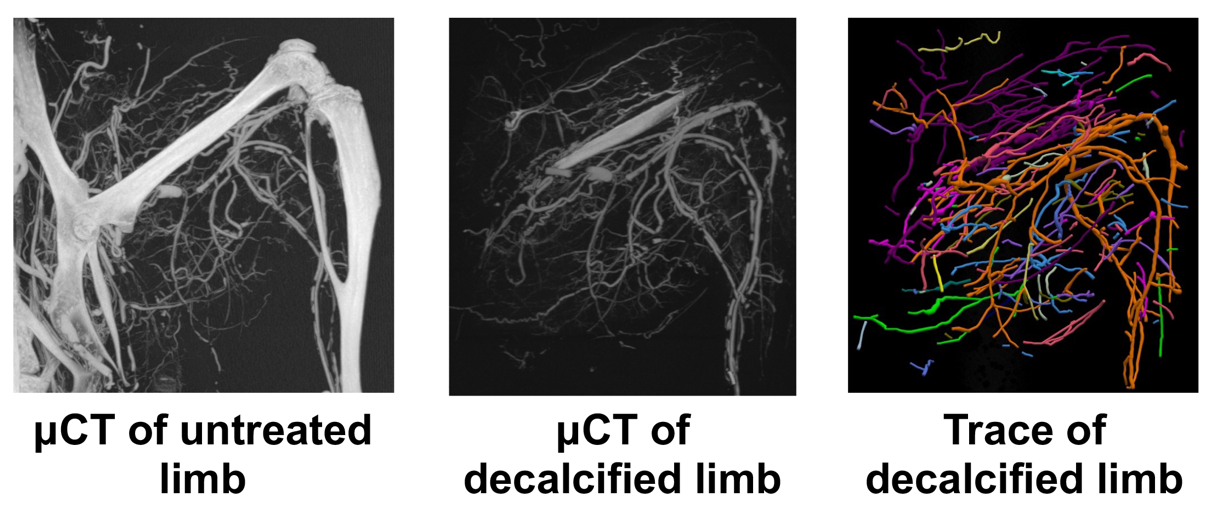

**Supplemental Figure 3: Example of decalcification process used to image polymer-filled vessels in the hindlimb.**
